# One-pot virus detection based on isothermal amplification coupled with temperature-activated argonaute

**DOI:** 10.1101/2023.10.16.562499

**Authors:** Xingyu Ye, Zhonglei Li, Zhengwei Fang, Nan Shen, Yuanjie Zhou, Peng Sun, Heshan Xu, Qian Liu, Yan Feng

## Abstract

Advances in programmable nucleases like CRISPR-associated protein (Cas) and Argonaute (Ago), combined with isothermal amplification, have made point-of-care testing (POCT) more accessible. However, the specific binding of the nuclease resulted in compatibility issues between the amplification and nuclease systems, substantially limiting the feasibility of a one-step workflow. Here, a temperature control solution based on immobilized thermotolerant *Pyrococcus furiosus* Ago (*Pf*Ago) has been proposed. The use of immobilized *Pf*Ago can effectively prevent interference with loop-mediated isothermal amplification (LAMP) at 65°C and accurately identifies amplicons when activated at 95°C. Following optimization, a sensitivity of 0.6 copies/μL was achieved within 45 minutes, and high specificity was verified with no cross-reactivity among 22 other viruses. Additionally, the multiplex detection was designed for herpes virus sensing, with agreements of 86.4% for positive and 100% for negative samples. Our research presents an effective method for combining amplification and cleavage through the use of controllable nucleases, significantly improving the clinical applicability of diagnostic techniques dependent on programmable nucleases.

## Introduction

To facilitate timely responses to pandemic outbreaks, a point-of-care test (POCT) technology with high sensitivity is required. However, current nucleic acid test systems performed in central laboratories have exposed issues such as high transportation costs, cumbersome sample storage, prolonged turnaround time, and expertise requirements (1-3). Isothermal nucleic acid amplification (INA) technologies such as loop-mediated isothermal amplification (LAMP) and recombinase polymerase amplification (RPA) have enabled the development of easy-to-use POCTs that do not demand specialized training or large-scale equipment (4-7). Yet, INAs have some limitations, e.g. low precision and restrictions in multiplex detection (8-13). Nevertheless, the ease of use of INAs provides a valuable diagnostic tool for pandemic response.

Programmable endonuclease-based biosensors have emerged as a promising approach for nucleic acid detection, particularly for reducing false positives in isothermal amplification reactions. CRISPR associated endonucleases, such as Cas12 and Cas13, are effective due to their sequence-dependent trans-cleavage activity triggered by specific amplicons, producing reliable results for nucleic acid testing (14-18). However, the necessity for separate INA and Cas systems may result in contamination and complex operations. Therefore, there is an increasing interest in developing an one-pot detection system. To integrate the Cas cleavage and INA system, various approaches have been explored, such as optimization of components (19-25), microfluidic device (26-29), modification/blocking of crRNA (30-34), engineering of Cas protein (35) and physical isolation (36-42). However, most of these methods rely on unique consumables and instruments, limiting their practicality. Moreover, further validation is needed to confirm the universality of target sequence recognition..

Argonaute proteins (Agos) have emerged as a promising alternative to CRISPR/Cas enzymes due to their ssDNA-directed programmable nuclease activity (43). One notable advantage of using Agos is their orthogonality, as it allows for the detection of multiple targets using a single enzyme without the need for a PAM or PFS recognition sequence (44-46). This makes Agos particularly suitable for multiplex detection applications, effectively reducing the number of enzymes required for sensing multiple targets (47, 48). In contrast, Cas12/13’s trans-cleavage activity requires the use of multiple enzymes for sensing multiple targets, thus necessitating further optimization of the CRISPR components to improve their applicability. Hence, Agos represent a promising nuclease tool for precise and multiplex detection. In general, Thermophilic Agos exhibit distinct sensing capabilities for dsDNA cleavage, which can be utilized in combination with PCR or INA techniques to enable virus detection (49-51). However, most Ago-based methods still require a two-step reaction due to limitations in the integration of amplification and cleavage systems. Although two recently systems, named Scalable and Portable Testing (SPOT) and Multiplexed Argonaute-based nucleic acid detection (MULAN), as primarily one-pot reactions, further specialized consumables and instruments are still necessary (51, 52).

In this study, we developed a One-Pot Test on IMmobilized Argonaute and LAMP, called as OPTIMAL, for detecting SARS-CoV-2 and classifying herpesviruses. To enhance LAMP efficiency, *Pf*Ago’s binding and cleaving capabilities were blocked by immobilization. Following LAMP amplification, the system is activated via heating to initiate *Pf*Ago sensing. The combination of all components within a single tube creates a closed and portable system. Our findings demonstrate that the detection system is highly effective, achieving a sensitivity of 0.6 copies/μL with no cross-reactivity among 22 other viruses. Finally, clinical validation of herpesvirus classification confirms the potential usefulness of this method for multiplex detection in clinical practice.

## Results

### Design of OPTIMAL via immobilized *Pf*Ago

It is commonly knowledge that Agos initially binds to nucleic acids before undergoing specific cleavage. However, to effectively carry out a one-pot reaction, the binding function of *Pf*Ago must first be inhibited to increase the availability of targets for amplification. At the optimal temperature of LAMP reaction (65°C), *Pf*Ago exhibits no cleavage activity but strongly binds to various nucleic acid fragments, including ssDNA and ssDNA-reporter dsDNA complex (Fig. 1A). As a result, the addition of *Pf*Ago to the reaction significantly hinders LAMP amplification even when using a high template concentration of 3000 copies/μL (Fig. 1B). Moreover, a low concentration of *Pf*Ago does not initiate the stepwise cleavage of amplicons and reporters (Fig. 1C). Due to the importance of developing one-pot systems for point-of-care testing (POCT) applications, we aimed to devise a method that easily modulates *Pf*Ago’s binding activity.

**Fig. 1.**
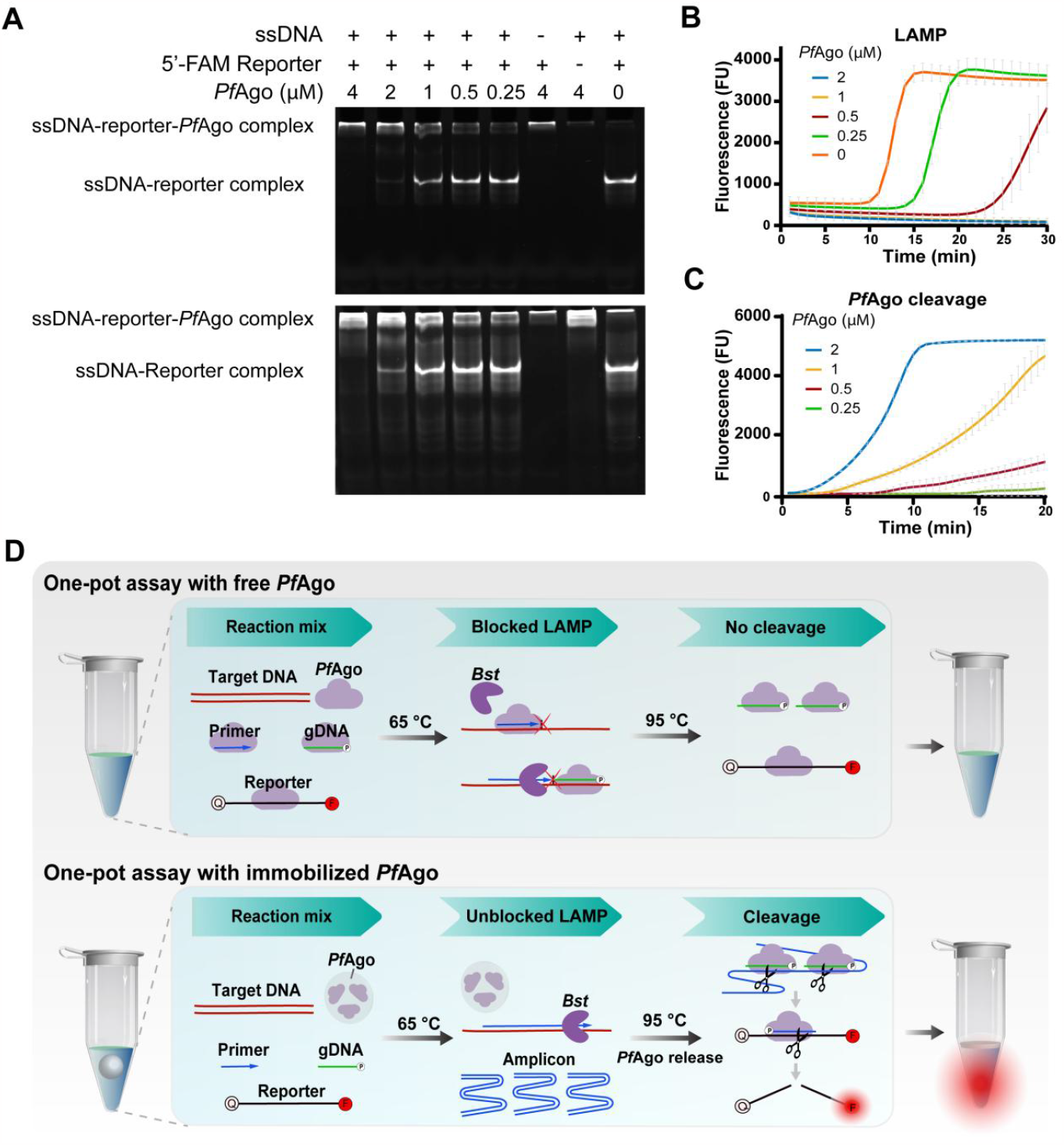
Design and strategy of immobilized *Pf*Ago-based detection. (A) Verification of *Pf*Ago’s ability to bind ssDNA, reporter and ssDNA-reporter complex using an electrophoresis mobility shift assay (EMSA). Reporter and ssDNA are inversely complementary and anneal to dsDNA. The upper gel was photographed before staining, and the bottom gel was photographed after staining. (B) LAMP amplification curve with *Pf*Ago added. (C) *Pf*Ago cleavage of reporters triggered by LAMP amplificons. (D) Principle of immobilization and thermo-controllable release of *Pf*Ago. (E) Schematic of immobilized *Pf*Ago-based one-pot detection. Data are represented as mean ± standard deviation (s.d.) (n = 3 replicates).

A one-pot procedure was developed to activate *Pf*Ago by taking advantage of the different reaction temperatures for LAMP and *Pf*Ago. This low-cost and efficient approach was achieved through solid immobilization, which is a suitable method for large-scale preparation. It was found that *Pf*Ago exhibited excellent thermostability over 99 °C (Fig. S1), which allowed it to withstand the solid immobilization process. Initially, the immobilization with solid paraffin was used to block *Pf*Ago at a temperature below the melting point. During the reaction, when the temperature exceeded the melting point of the solid paraffin, *Pf*Ago became active and flowed into the aqueous solution due to its hydrophilicity. Moreover, the melted paraffin floated on the surface of the reaction solution, preventing evaporation and contamination during the entire procedure.

By utilizing immobilized *Pf*Ago, we present a novel one-pot detection method named OPTIMAL (One-Pot Test on IMmobilized Argonaute and LAMP) that allows for efficient blocking of *Pf*Ago during the LAMP reaction, while enabling multiplex detection with improved sensitivity (Fig. 1C). In this one-pot reaction, amplification and cleavage occur sequentially with increasing temperature. The LAMP amplification occurs initially at 65°C without interference from immobilized PfAgo. Then, at 95°C, *Pf*Ago is released to cleave the amplicon guided by two specific gDNAs to generate renewed gDNA that initiates secondary cleavage of the reporters, resulting in fluorescence readout. The OPTIMAL method effectively overcomes the limitations of conventional two-step *Pf*Ago-based assays, offering a viable solution to address contamination and operational intricacies.

### Assessment of immobilized *Pf*Ago cleavage system

To evaluate the effectiveness of immobilized *Pf*Ago, we tested immobilized *Pf*Ago beads of varying diameters for single-stranded DNA cleavage, showing that the beads with a diameter of 2 mm exhibited the most effective and stable activity (Fig. 2A). These beads were then incorporated into the LAMP reaction, and did not significantly impact amplification efficiency (Fig. 2B-D). To assess the durability of immobilized *Pf*Ago under normal environmental conditions, we tested it at room temperature and 2-8°C over 30 days. We found that the immobilized *Pf*Ago remained active throughout, implying safe and damage-free storage and shipping (Fig. 2E). By optimizing the *Pf*Ago reaction for compatibility with the magnesium ion (Mg^2+^) activated LAMP system, we determined that 8 mM of Mg^2+^ were the ideal divalent metal ions for the immobilized *Pf*Ago cleavage reaction. As expected, under these optimized Mg^2+^ concentrations, the immobilized *Pf*Ago retained over 90% activity compared to typical manganese ion (Mn^2+^) activated cleavage (Fig. 2F).

**Fig. 2.**
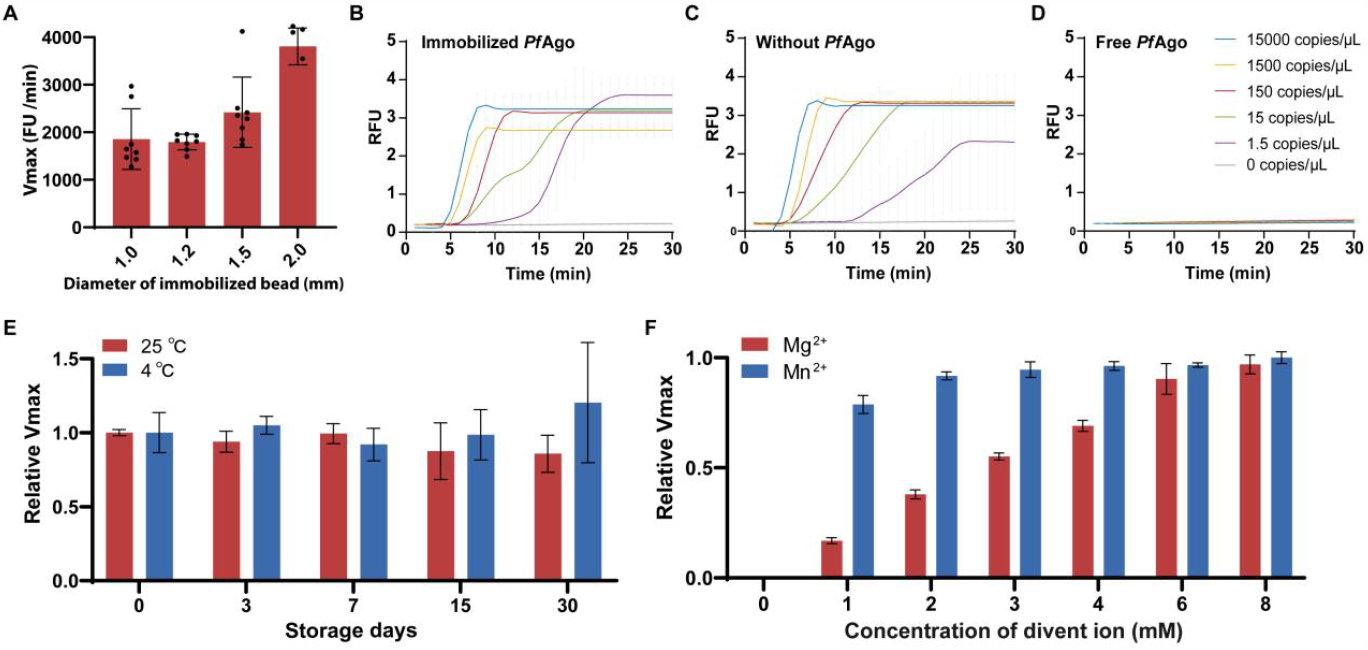
Assessment of immobilized *Pf*Ago cleavage system. (A) Activity of immobilized *Pf*Ago beads with diverse diameters. Vmax represents the maximum velocity of fluorescent unit growth from ssDNA reporter cleavage. LAMP amplification curve with immobilized *Pf*Ago (B), without *Pf*Ago (C) and with free *Pf*Ago (D) using gradient loads of templates. The RFU on the vertical axis represents the normalized relative fluorescence units. (E) Validation of storage time of immobilized *Pf*Ago at room temperature (RT). The relative Vmax represents the ratio to the Vmax at the timepoint of day 0. (F) Optimization of divalent metal ion and buffer of *Pf*Ago cleavage system. The relative Vmax represents the ratio to the Vmax of 8 mM Mn^2+^. Data are represented as mean ± standard deviation (s.d.) (n = 3 replicates).

### Establishment and evaluation of one-pot detection system

During LAMP, Mg^2+^ binds with pyrophosphoric acid, forming a pyrophosphate by-product that can reduce the effectiveness of subsequent *Pf*Ago reactions. To address this issue, we added pyrophosphatase (PPase) to prevent the formation of magnesium pyrophosphate precipitation (Fig. 3A). We used SARS-CoV-2 as detection object, and designed gDNAs and reporters targeting N gene (Fig. S2). The results demonstrated that one unit of PPase was enough to prevent the Mg2+ consumption (Fig. 3B), while increasing immobilized *Pf*Ago to three beads obtained optimized efficiency without sacrificing sensitivity (Fig. 3C). Finally, reporter and gDNA concentrations were optimized for shorter amplification and cleavage time (Fig. S3 and S4).

**Fig. 3.**
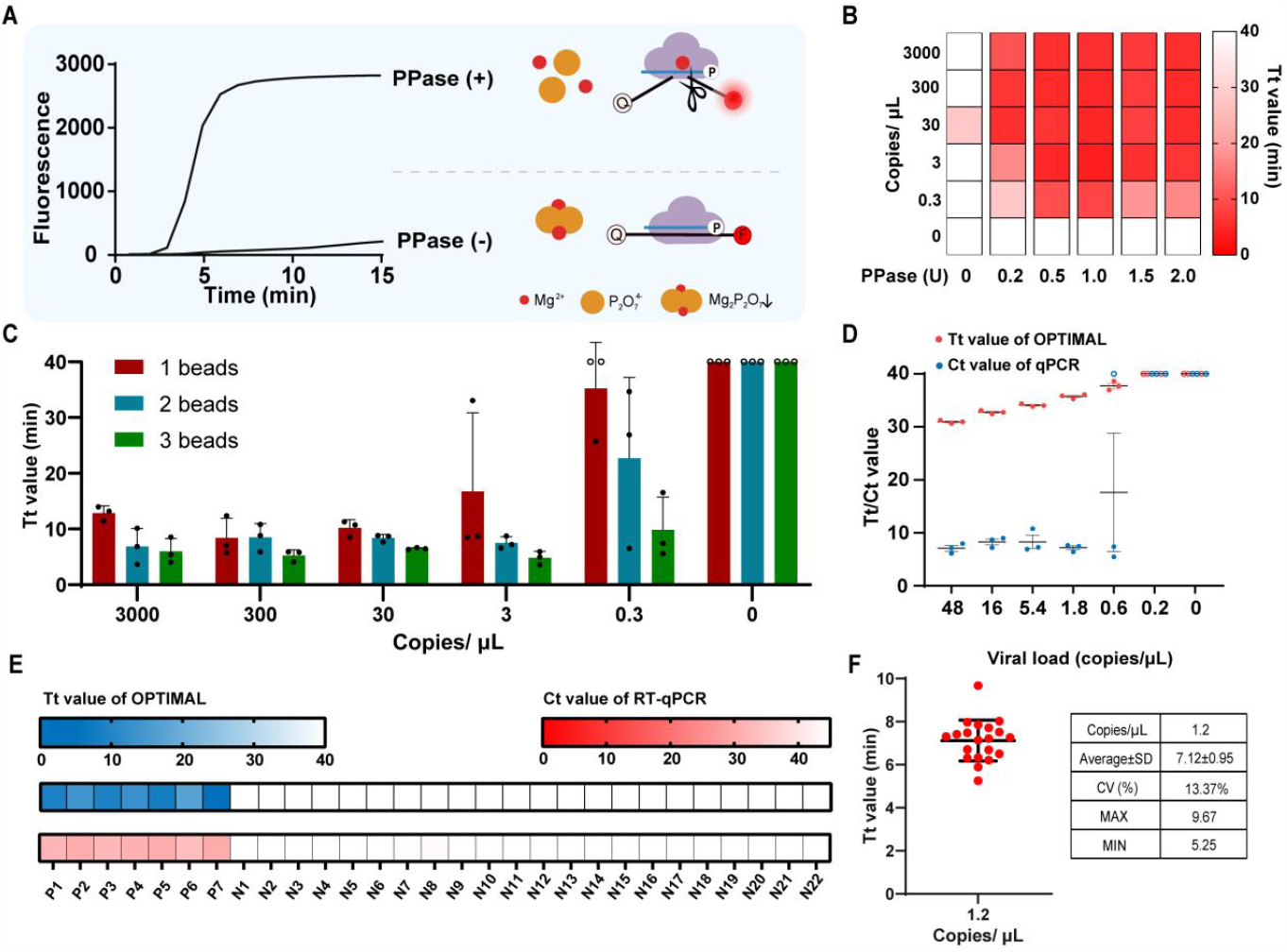
Optimization and evaluation of OPTIMAL. (A) Schematic of one-pot detection with PPase added to maintain Mg concentration. (B) Optimization of PPase concentration and (C) amount of immobilized PfAgo beads for one-pot detection. The circles indicate no determined value within 40-min reaction. (D) Evaluation of limit of detection using inactivated SARS-CoV-2. The circles indicate no determined Tt/Ct value within 40-min/40-cycle reaction. (E) Specificity assay wherein the cross-reactivity for 28 common respiratory pathogens was tested. P1: Wild strain of SARS-CoV-2, P2: Beta strain of SARS-CoV-2, P3: Gamma strain of SARS-CoV-2, P4: Delta strain of SARS-CoV-2, P5: Omicron strain of SARS-CoV-2, P6: Wild strain of SARS-CoV-2, P7: N gene of wild SARS-CoV-2, N1: Legionella pneumophila, N2: Klebsiella pneumoniae, N3: Streptococcus pneumoniae, N4: Haemophilus influenzae, N5: Adenovirus type 3, N6: Mycoplasma pneumoniae, N7: Chlamydia pneumoniae, N8: Parainfluenza type 1, N9: Respiratory Syncytial Virus type A, N10: Bordetella pertussis, N11: Human coronavirus OC43, N12: Human coronavirus NL63, N13: Human coronavirus HKU-1, N14: Human coronavirus 229E, N15: Influenza A virus subtype H7N9, N16: Influenza A virus subtype H5N1, N17: Influenza B victoria lineage, N18: Influenza A virus subtype H1N1(2009), N19: Influenza A virus subtype H3N2, N20: Epstein-Barr virus, N21: MERS-CoV (Pseudoviruses with partial genes), N22: Simulated negative swab samples. (F) 20 replicates of low-load SARS-CoV-2 were tested to evaluate the reproducibility. (G) The coefficient of variation index about the reproducibility assay. Data are represented as mean ± standard deviation (s.d.) (n = 3 replicates). The depth of red and blue in the heat map represents the average Tt value and Ct value of three replicates, respectively.

To evaluate the effectiveness of OPTIMAL, we conducted experiments to measure the limit of detection (LoD), specificity, and reproducibility. Using inactivated SARS-CoV-2 cultured in vitro as templates, we found that the LoD was 0.6 copies/μL (Fig. 3D), which corresponds to a mean Ct of 37.8 as determined by a commercial qPCR kit. Importantly, we discovered no cross-reactivity with 21 different pathogens, and all SARS-CoV-2 mutant strains were detected, thus confirming the high specificity (Fig. 3E). Finally, we tested 20 samples with a load twice the LoD, equaling 1.2 copies/μL, and achieved a coefficient of variation with only 13% for Tt values. These results were obtained in less than 10 minutes, highlighting the rapidity and reliability of OPTIMAL (Fig. 3F and 3G).

### Development of visual home-testing

For this one-pot visual detection, we developed a low-cost isothermal detection by a handheld device. We filtered a disposable microburette with high titration precision, instead of the laboratory’s pipettor (Fig. S5). Viral lysate was used to release viral RNA, and combined with a handheld detector, a home self-testing process for detecting SARS-CoV-2 was developed (Fig. 4A and S6). Reporters of varying lengths were designed to increase the signal-to-noise ratio (SNR) for endpoint detection (Fig. 4B and 4C). We found that an 18-nt reporter achieved the highest SNR and did not significantly impact detection time. The simulated nasopharyngeal swab sensitivity was validated at 1.3 copies/uL, equivalent to a qPCR Ct value of 37.32 (Fig. 4D and 4E). The OPTIMAL detection time was reduced to 45 minutes, comprising a 30-minute amplification and 15-minute read-out, compared to over 60 minutes for qPCR.

**Fig. 4.**
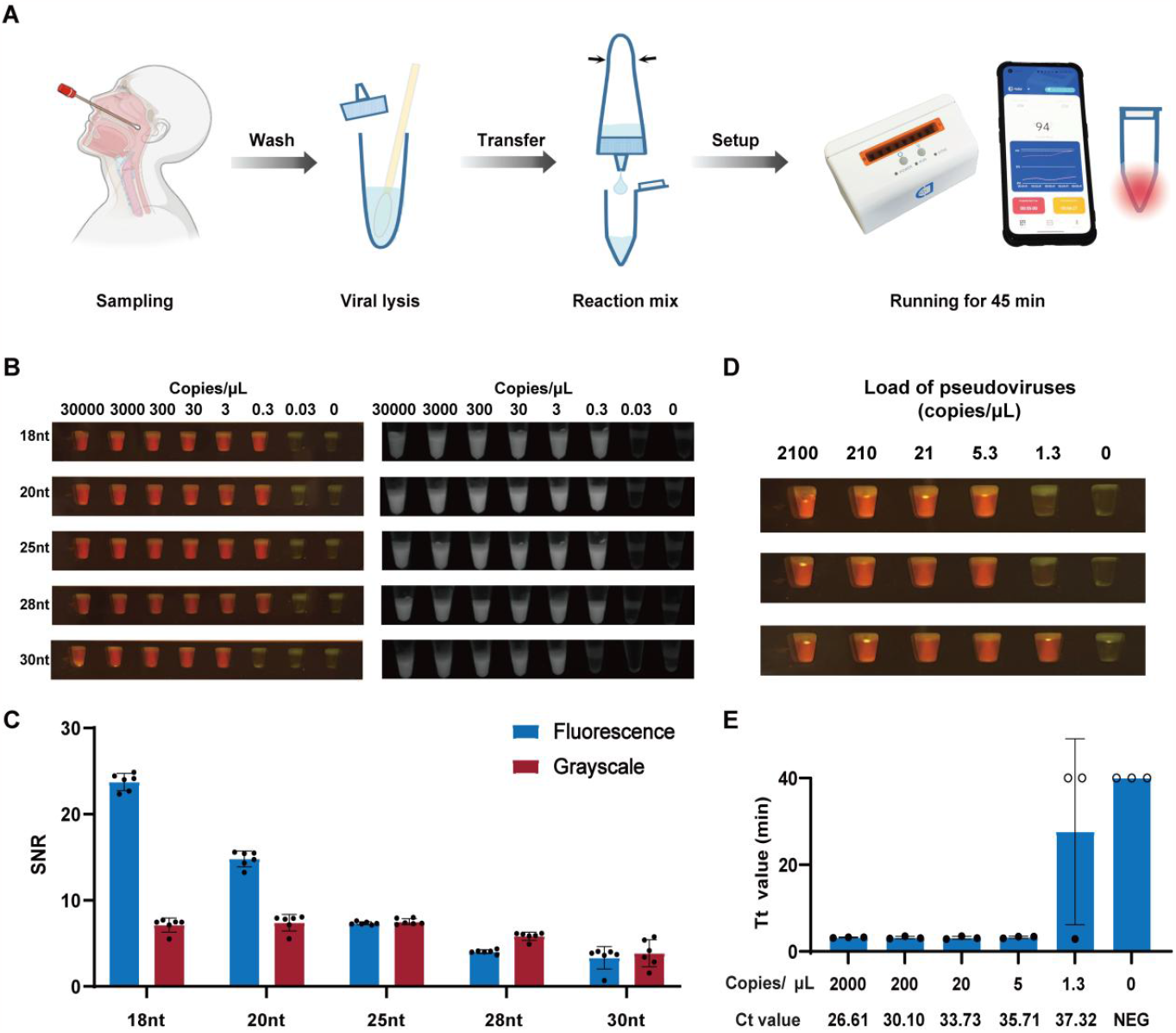
Development of visual home-testing. (A) The procedure of visual home-testing base on OPTIMAL. (B) Visual detection results using reporters of diverse length displayed in photogragh (left) and grayscale (right). (C) Signal-to-noise ratio (SNR) of fluorescence with reporters of diverse length. (D) Sensitivity assay of home-testing using pseudoviruses in throat swabs as samples. (E) The Tt value and sensitivity with fluorescent readout versus the Ct value of qPCR. The circles indicate no determined value within 40-min reaction. Data are represented as mean ± standard deviation (s.d.) (n = 3 replicates).

### Construction and clinical validation of multiplex detection

The human herpesviridae family, including cytomegalovirus (CMV), Epstein-Barr virus (EBV), and varicella-zoster virus (VZV), are prevalent DNA viruses with similar clinical manifestations. Rapid differentiation and diagnosis of these viruses in outpatient clinics are essential for guiding appropriate therapy. However, no POCT multiplex detection methods are currently available for these viruses. To address this gap, we have developed an OPTIMAL method that can simultaneously detect all three viruses in one tube without lid-opening and manual operation (Fig. 5A). For the single-plex detection, each virus can be detected with a sensitivity as low as 0.3 copies/μL (Fig. S7). After optimizing the primer ratio for multiplex detection (Fig. S8-10), we achieved a multiplex detection with a sensitivity of 3 copies/μL for each virus, without any cross-reactivity (Fig. 5B).

**Fig. 5.**
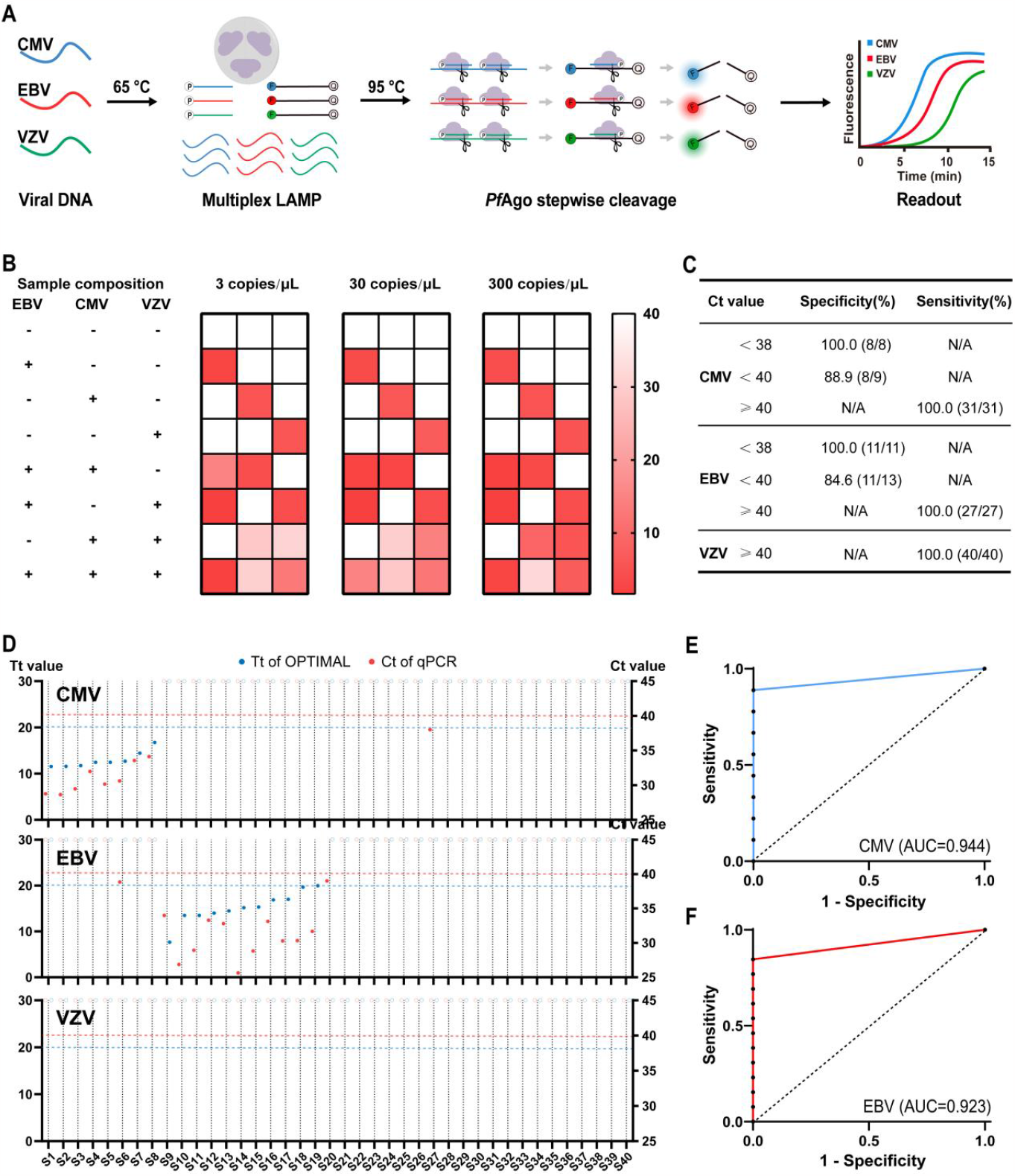
Multiplex detection based on OPTIMAL. (A) Schematic of multiplex detection with immobilized *Pf*Ago. (B) Triplex detection with mixed samples of EBV, CMV and VZV plasmids. (C) Sensitivity and specificity of triplex detection for clinical samples. (D) Comparison of OPTIMAL and commercial qPCR via a parallel test of clinical samples. The red dash lines represent the cut-off Ct value of 40 for qPCR and the red dash lines represent the cut-off Tt value of 20 for OPTIMAL. The circles indicate no determined value within 30-min reaction. (E) Receiver operating characteristic (ROC) curve of CMV detection. (F) Receiver operating characteristic (ROC) curve of EBV detection. The depth of red in the heat map represents the average fluorescence intensity of three replicates.

The optimized multiplex detection method was validated on 40 samples taken from children exhibiting related symptoms, and its performance was compared to that of a commercial qPCR kit. When the positive threshold was set at Ct<40, sensitivities of 88.9% (8/9) and 84.6% (11/13) were attained for CMV and EBV, respectively. When the positive threshold was set at Ct<38, all samples displayed 100% sensitivity and 100% specificity for each virus (Fig. 5C, Table S1). CMV and EBV were found to be the main causative pathogens, while VZV went undetected by either the OPTIMAL or qPCR approach (Fig. 5D). Using a Ct value of 40 as the cut-off value, the receiver operating characteristic (ROC) curve demonstrated that the area under the curve (AUC) were 0.944 and 0.923 for CMV and EBV respectively, confirming the excellent predictive accuracy of this multiplex OPTIMAL test (Fig. 5E, F). In summary, our findings illustrate the effectiveness of OPTIMAL for clinical practice.

## Discussion

The advancement of molecular diagnostics is attributed to innovative nucleases such as CRISPR/Cas and Agos, which provide highly sensitive and portable detection platforms (43, 53, 54). Agos exhibit specific cleavage activity across multiple targets, enabling the development of cost-effective and simple multiplex systems. However, commercializing detection technology based on Agos and CRISPR/Cas has faced challenges due to aerosol contamination and the manual two-step operation. To overcome these issues, immobilization provides a thermal controlled blocking of *Pf*Ago that is compatible with LAMP amplification. Compared to other blocking methods (30-34) and physical isolation (36-42), immobilization is more universal and economical, easy for large-scale preparation. In this study, immobilized *Pf*Ago was utilized in developing the OPTIMAL system, which features a one-pot detection strategy.

The OPTIMAL system demonstrated similar sensitivity to the two-step system, detecting as low as 0.6 copies/μL of SARS-CoV-2, outperforming the Cas12/Cas13 detection systems(55, 56). Additionally, OPTIMAL’s detection time was only 45 minutes, making it an attractive option for rapid response, paving a way for simpler, faster, and more accurate molecular diagnostic detection, particularly in the case of emerging viruses, such as SARS-CoV-2. Aided by handheld device, OPTIMAL enables versatile capabilities not only for one-pot detection but also for POCT and home testing. The device does not require a temperature cycling module or hot lid for OPTIMAL reaction, making it compact and cost-effective for timely prevention and control of future epidemics. Regarding the differential diagnosis of viral infections, OPTIMAL is superior in multiplex detection of herpesviruses, accurately differentiating the three viruses: VZV, EBV, and CMV, at low viral loads. In clinical validation, OPTIMAL demonstrates high agreement with commercial kits, with only a few positive samples going undetected at a Ct value above 38. This improvement ensures rapid differential diagnosis in practical applications.

Overall, OPTIMAL provides an optimal solution for one-pot, portable, multiplex programmable nuclease-based detection. It takes full advantage of *Pf*Ago’s biochemical features, including thermostability, precise cleavage, and multiplexity. The one-step strategy of OPTIMAL can also be integrated with other thermostable agos, such as *Fp*Ago, *Mf*Ago, as well as a range of isothermal amplification technologies, including RPA, EXPAR. This integration of immobilized agos in nucleic acid detection technology promotes the development of powerful molecular diagnosis, and enables timely epidemic prevention.

## Materials and methods

### Materials used in the experiments

All oligonucleotides and plasmids were purchased from Sangon Biotech and are listed in Table S2-6. Plasmids were quantified using the PicoGreen dsDNA Quantitation Kit (Life iLab Biotech). *Pf*Ago was expressed and purified as described previously (47). The solid immobilization process was jointly developed with Jiaohong Biotechnology Co., LTD. Bst DNA/RNA Polymerase (4.0) was used for LAMP, and thermostable pyrophosphatase (PPase) for reserving free Mg^2+^ was purchased from HaiGene Biotech. The QIAamp Viral RNA Mini Kit (Qiagen) was used for SARS-CoV-2 RNA extraction. A viral DNA extraction kit (Tiangen) was used for CMV, EBV, and VZV DNA extraction. Commercial RT-qPCR kits for viruses used in this study were provided by BioPerfectus.

### *Pf*Ago cleavage of ssDNA reporter

The reaction mix was prepared with the following components: 2.5 μL of 10× reaction buffer (150 mM Tris/HCl, 2.5 M NaCl, pH 8.0), 4 μM gDNAs, 2 μM ssDNA reporters, 0.4 mM MnCl_2_, and 1 μM *Pf*Ago, and then made up to 25 μL with nuclease-free water. The cleavage reactions were performed at 95°C for 30 min and the fluorescence was recorded using SLAN-96P (Hongshi Sci Tech).

### Electrophoresis mobility shift assays (EMSAs)

*Pf*Ago with various concentration (0.25, 0.5, 1, 2, 4 μM) was pre-incubated with 2 μM FAM labeled 30 nt-reporters and 2 μM 90 nt-ssDNA complementary to the 30 nt-reporters for 5 min at 65 °C in a typical LAMP reaction buffer (20 mM Tris-HCl, pH 8.8, 10 mM KCl, 8 mM MgSO4, 10 mM (NH4)2SO4, 0.1% Tween 20). Next, the samples were mixed with 10x EMSA loading buffer (0.1% bromophenol blue and 50% glycerin) and analyzed using 8% non-denaturing polyacrylamide gels. Subsequently, the unstained gels were visualized under blue light, and the GelRed-stained gels were visualized under ultraviolet light with a wavelength of 305 mm in a gel imaging system (Tanon Life Science).

### LAMP (RT-LAMP) assays

The primers for each virus were designed using Primer Explorer V5 (https://primerexplorer.jp/e/). The primer mix was prepared with defined concentrations of 1.6 μM internal primers (FIP/BIP), 0.2 μM external primers (F3/B3), 0.4 μM loop primers (LF/LB), 8 U Bst 4.0 DNA/RNA Polymerase, 1× Bst polymerase buffer 6 mM MgSO_4_, 0.5 μL of SYTO-9 dsDNA dye (NEB BioLabs, for LAMP fluorescent curve) was used to monitor the real-time amplification signals. DNA or RNA templates (5 μL) were added, and nuclease-free water was added to a total volume of 25 μL per reaction. reaction was performed at 65 °C for 30 min.

### *Pf*Ago cleavage assays

For the *Pf*Ago cleavage of LAMP products, the master mix was prepared with the following components: 2.5 μL of 10× reaction buffer (150 mM Tris/HCl, 2.5 M NaCl, pH 8.0), 2.5 μM gDNAs, 0.5 μM reporters, 0.4 mM MnCl_2_, and 10 μM *Pf*Ago, and then made up to 25 μL with nuclease-free water. The master mix was mixed with 25 μL of the LAMP products.

The cleavage reactions were performed at 95 °C for 30 min, and the real-time fluorescence was measured using SLAN-96P (Hongshi Sci Tech).

### One-pot assays integrating LAMP and *Pf*Ago cleavage for SARS-CoV-2 detection

The primer mix, Bst polymerase, and Bst buffer were prepared as described in the aforementioned LAMP assays. Other components were prepared with defined concentrations 2 μM N-gene gDNAs, 0.4 μM ROX-labeled N-gene reporters, 8 mM MgSO_4_, three beads of immobilized *Pf*Ago and 1 U PPase. The reactions were performed at 65 °C for 30 min followed by 95°C for 15-30 min, and the real-time fluorescence was measured using SLAN-96P (Hongshi Sci Tech).

### Home-testing assays for SARS-CoV-2 detection

For the home-testing assays for SARA-CoV-2 detection, the primer mix was prepared at defined concentrations of 32 μM FIP and BIP, 4 μM F3 and B3, 8 μM LF and LB. The premix was set in 40 μL volume as the one-pot assay with the following components: 3.75 μL of primer mix, 7.5 μL of 10× Bst Buffer, 3 μL of 8 U Bst 4.2 DNA/RNA Polymerase, 4.5 μL of 100 mM Mg+, 10.5 μL of 10 mM dNTP mixture, 1.5 μL of 100 μM N-gene gDNAs, 6 μL of 10 μM ROX-labeled N-gene reporters, three beads of immobilized *Pf*Ago and 1.5 U PPase. Then one drop of swab lysates or references with a volume of approximately 35 uL was added to the premix. The reactions were performed at 65 °C for 30 min followed by 95°C for 15-30 min, and the endpoint fluorescence was observed using WeD (Hangzhou Institute for Advanced Study, UCAS).

### Multiplex OPTIMAL assays for the detection of CMV, EBV and VZV

For multiplex detection, the primer mix was prepared for each virus at defined concentrations of 32 μM FIP and BIP, 4 μM F3 and B3, 8 μM LF and LB. The reaction mix was set in 25 μL volume as the one-pot assay mentioned above, except for the following components: 0.8 μL of CMV primer mix, 0.4 μL of EBV primer mix, 0.4 μL of VZV primer mix, 2 μM gDNA specific for each virus, 0.4 μM of FAM-labeled reporter for CMV, ROX-labeled reporter for EBV and VIC-labeled reporter for VZV. The reactions were performed at 65 °C for 30 min followed by 95°C for 30 min, and the real-time fluorescence was measured using SLAN-96P (Hongshi Sci Tech).

### Devices and consumables used in the home-testing

The handheld device was generously provided by Prof. Jinzhao Song’s group from the Cancer Hospital of the University of Chinese Academy of Sciences (Zhejiang Cancer Hospital), Institute of Basic Medicine and Cancer (IBMC), Chinese Academy of Sciences. The determined disposable microburettes were purchased from Zhejiang Aicor Medical Technology Co., Ltd, and viral lysates were purchased from Wuhan Hzymes Technology Co., Ltd.

### Acquisition of clinical samples and references

Serum samples for herpes virus detection were collected from the Shanghai Children’s Medical Center. The SARS-CoV-2 pseudoviruses were purchased from Co-Bioer Biosciences. The *in vitro*-cultured respiratory pathogens were purchased from the National Institute for Food and Drug Control of China.

### Credit authorship contribution statement

**Xingyu Ye:** Methodology, Conceptualization, Investigation, Writing - original draft, Writing - review & editing, Visualization. **Zhonglei Li:** Methodology, Conceptualization, Validation. **Zhengwei Fang:** Methodology, Visualization, Writing - original draft, Formal analysis. **Nan Shen:** Investigation, Sample preparation, Validation. **Yuanjie Zhou:** Investigation, Sample preparation, Validatio.. **Peng Sun:** Methodology, Visualization, Formal analysis. **Heshan Xu:** Methodology. **Qian Liu:** Writing - original draft, Writing - review & editing, Visualization, Funding acquisition, Supervision. **Yan Feng:** Supervision, Conceptualization, Writing - review & editing, Funding acquisition.

## Declarations of competing interests

Xingyu Ye, Zhonglei Li, Zhengwei Fang, Qian Liu and Yan Feng are coinventors on a patent application related to this manuscript (application number, CN202311154019.7, applied September 8, 2023). The remaining authors declare no conflicts of interest.

## Acknowledgments

This work was supported by the Ministry of Science and Technology (2020YFA0907700), National Natural Science Foundation of China (31770078), Shanghai Pilot Program for Basic Research-Shanghai Jiao Tong University (21TQ1400204), and National Key Research & Development Program of China (2018YFA0900403). We would also like to express our sincere appreciation to Prof. Jinzhao Song from the Cancer Hospital of the University of Chinese Academy of Sciences (Zhejiang Cancer Hospital), Institute of Basic Medicine and Cancer (IBMC), Chinese Academy of Sciences for his invaluable insights and constructive discussions throughout the development of our visual home-testing method.

## Appendix A. Supplementary data

The following is the Supplementary data to this article:

## Supplementary Information

