## Supplemental Information for "One-pot virus detection based on isothermal amplification coupled with temperature-activated argonaute"

Supplementary Information

### Temperature-controlled activation enables one-pot thermophilic Argonaute-based diagnostics

*Address correspondence to:
Prof. Yan Feng, MD
Biochemistry School of Life Sciences and Biotechnology
Shanghai Jiao Tong University
800 Dongchuan Rd., Shanghai 200240, China


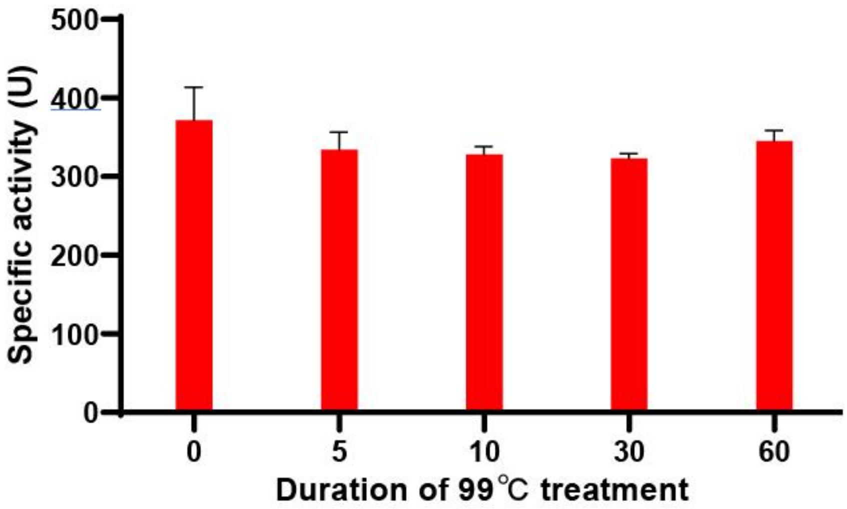


**Fig. S1.** Assessment of the specific activity of lyophilized *Pf*Ago after 99 °C treatment. Data are represented as mean ± standard deviation (s.d.) (n = 3 replicates).

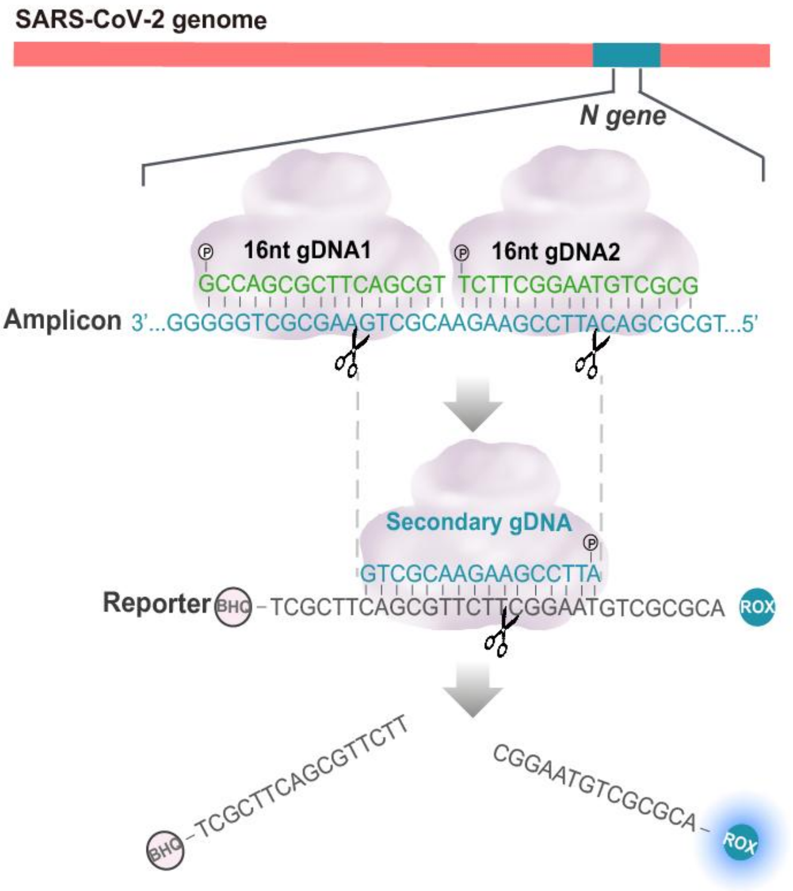


**Fig. S2.** Design of gDNA and reporter for the Argonaute-based detection of SARS-CoV-2 N gene. The purple cloud represents the *Pf*Ago.

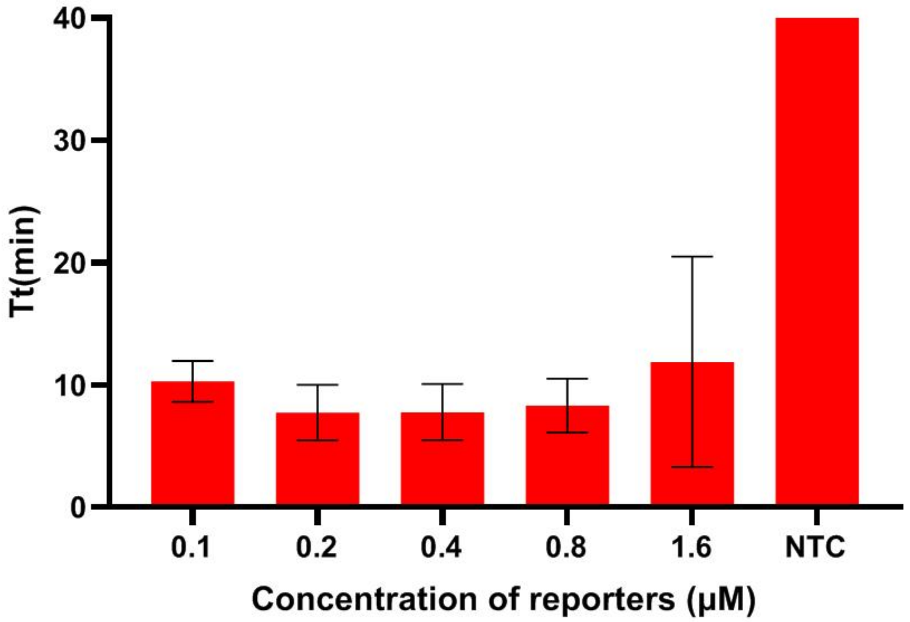


**Fig. S3.** Optimization of reporter concentration used for the OPTIMAL detection of SARS-CoV-2 N gene. The Tt value represents the time to threshold. Data are represented as mean ± standard deviation (s.d.) (n = 3 replicates).

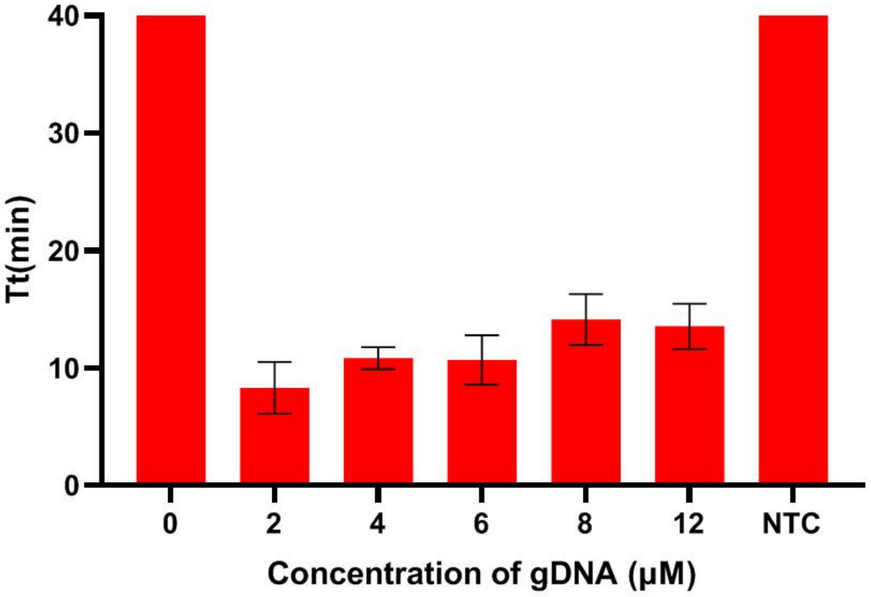


**Fig. S4.** Optimization of gDNA concentration used for the OPTIMAL detection of SARS-CoV-2 N gene. The Tt value represents the time to threshold. Data are represented as mean ± standard deviation (s.d.) (n = 3 replicates).

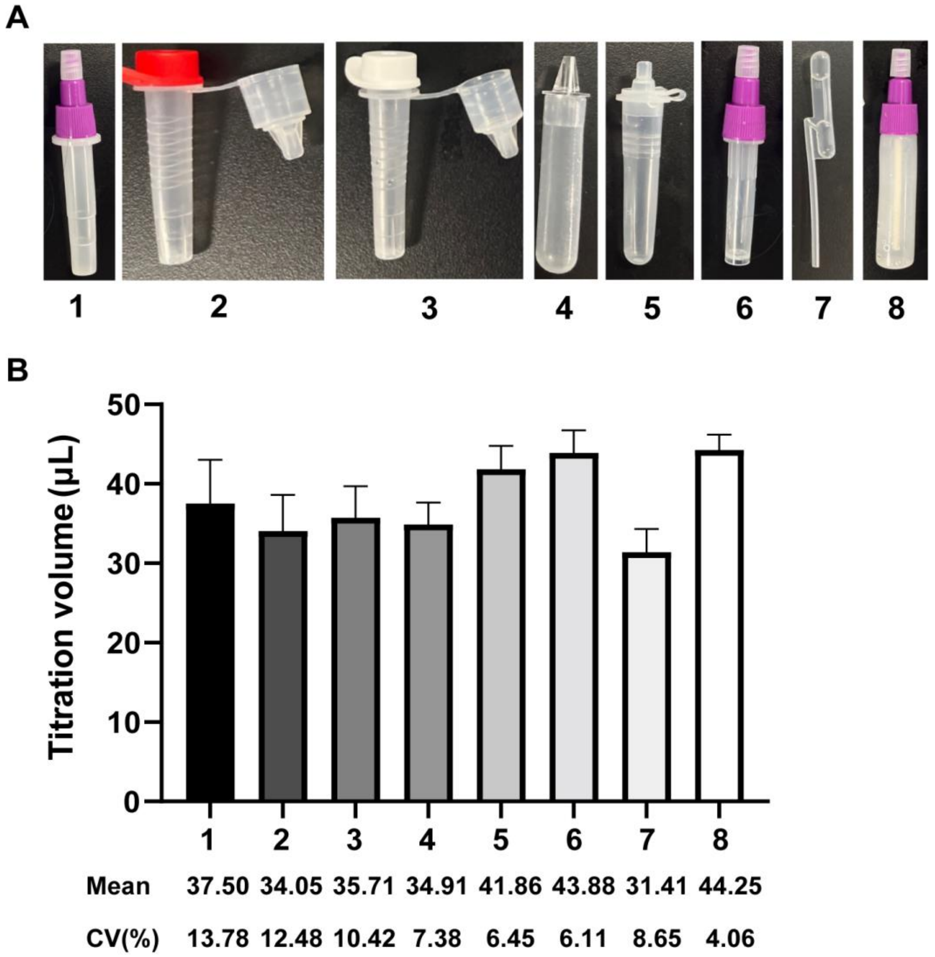


**Fig. S5.** Screening of disposable microburettes for precisely titration of viral lysates. (A) Candidates of disposable microburettes used for screening. (B) Evaluation of precision for the titration volume of disposable microburettes. No. 4 microburettes was used for the home-testing assays. Data are represented as mean ± standard deviation (s.d.) (n = 8 replicates).

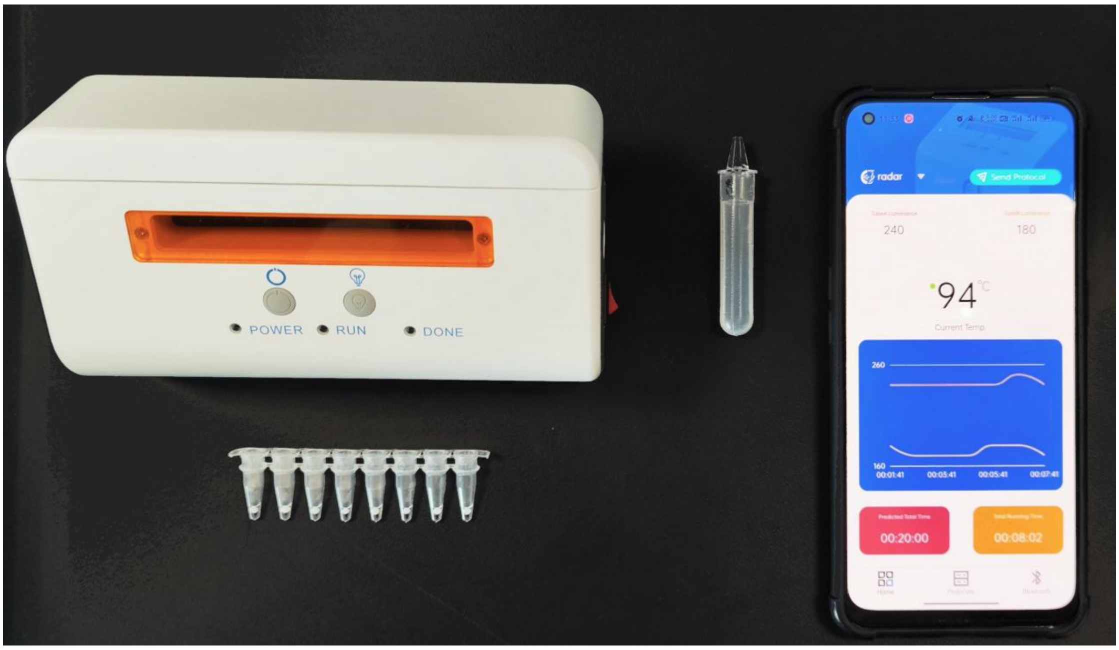


**Fig. S6.** The handheld device, consumables and presetting OPTIMAL reagents used for home-testing. The phone can display the temperature status and transmit the test results.

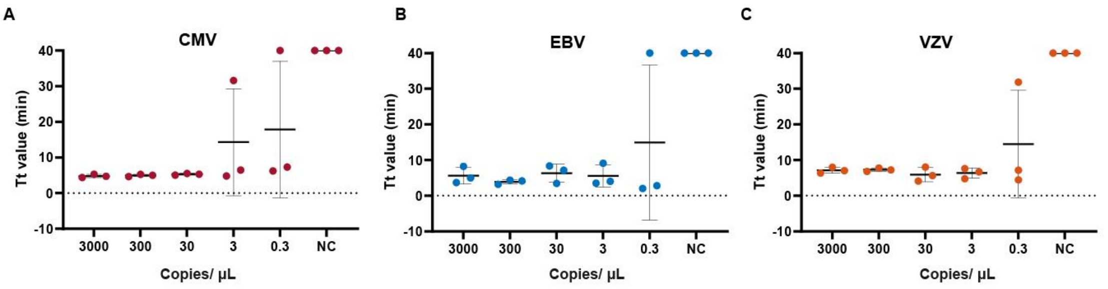


**Fig. S7.** Assessment of LoD for the single-plex detection of CMV (A), EBV (B) and VZV (C), respectively. Data are represented as mean ± standard deviation (s.d.) (n = 3 replicates).

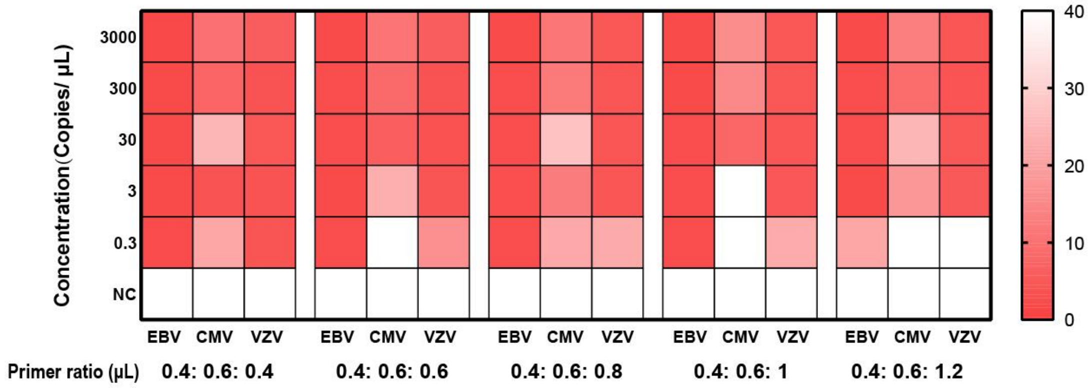


**Fig. S8.** Optimization of VZV primer ratio to others for multiplex detection of EBV, CMV and VZV. The amount of EBV and CMV primer mix were remained at 0.4 μL and 0.6 μL. Data are represented as mean ± standard deviation (s.d.) (n = 3 replicates).

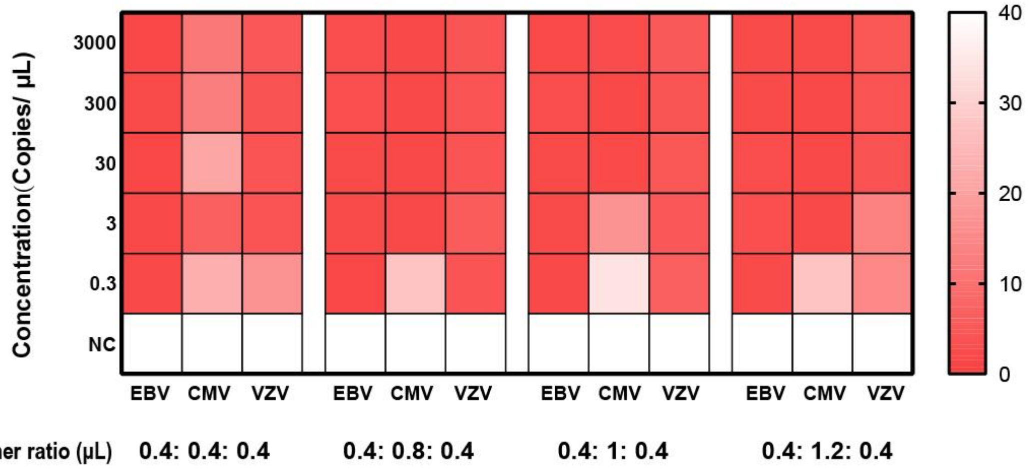


**Fig. S9.** Optimization of CMV primer ratio to others for multiplex detection of EBV, CMV and VZV. The amount of EBV and VZV primer mix were remained at 0.4 μL and 0.4 μL. Data are represented as mean ± standard deviation (s.d.) (n = 3 replicates).

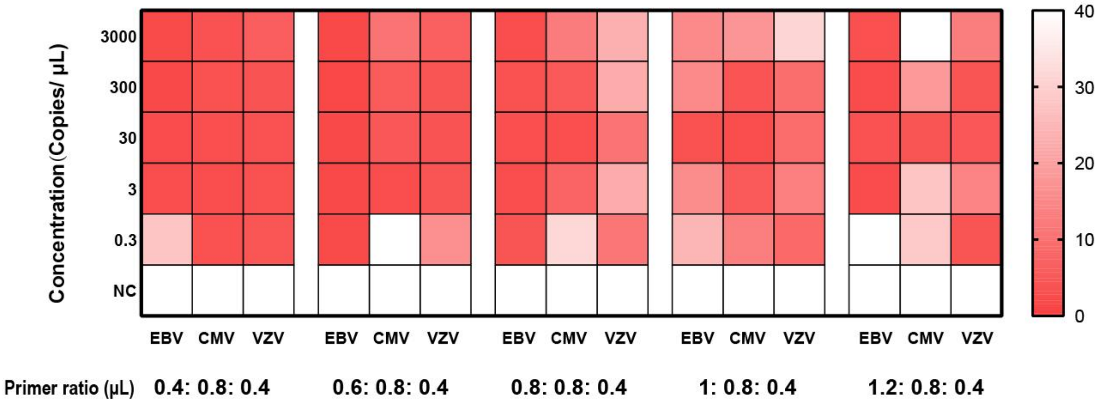


**Fig. S10.** Optimization of EBV primer ratio to others for multiplex detection of EBV, CMV and VZV. The amount of CMV and VZV primer mix were remained at 0.8 μL and 0.4 μL. Data are represented as mean ± standard deviation (s.d.) (n = 3 replicates).

| Samples | CMV | | EBV | | VZV | |
|---|---|---|---|---|---|---|
| | Ct value | Tt value | Ct value | Tt value | Ct value | Tt value |
| **S1** | 28.76 | 11.56 | - | - | - | - |
| **S2** | 28.64 | 11.59 | - | - | - | - |
| **S3** | 29.46 | 11.75 | - | - | - | - |
| **S4** | 31.97 | 12.45 | - | - | - | - |
| **S5** | 30.16 | 12.45 | - | - | - | - |
| **S6** | 30.62 | 12.69 | 38.84 | - | - | - |
| **S7** | 33.57 | 14.42 | - | - | - | - |
| **S8** | 34.12 | 16.75 | - | - | - | - |
| **S9** | - | - | 34.00 | 7.64 | - | - |
| **S10** | - | - | 26.87 | 13.51 | - | - |
| **S11** | - | - | 28.92 | 13.51 | - | - |
| **S12** | - | - | 33.30 | 13.98 | - | - |
| **S13** | - | - | 32.80 | 14.46 | - | - |
| **S14** | - | - | 25.63 | 15.17 | - | - |
| **S15** | - | - | 28.82 | 15.32 | - | - |
| **S16** | - | - | 33.13 | 16.88 | - | - |
| **S17** | - | - | 30.28 | 17 | - | - |
| **S18** | - | - | 30.34 | 19.65 | - | - |
| **S19** | - | - | 31.68 | 19.97 | - | - |
| **S20** | - | - | 39.00 | 7.64 | - | - |
| **S21** | - | - | - | - | - | - |
| **S22** | - | - | - | - | - | - |
| **S23** | - | - | - | - | - | - |
| **S24** | - | - | - | - | - | - |
| **S25** | - | - | - | - | - | - |
| **S26** | - | - | - | - | - | - |
| **S27** | 38.00 | - | - | - | - | - |
| **S28** | - | - | - | - | - | - |
| **S29** | - | - | - | - | - | - |
| **S30** | - | - | - | - | - | - |
| **S31** | - | - | - | - | - | - |
| **S32** | - | - | - | - | - | - |
| **S33** | - | - | - | - | - | - |
| **S34** | - | - | - | - | - | - |
| **S35** | - | - | - | - | - | - |
| **S36** | - | - | - | - | - | - |
| **S37** | - | - | - | - | - | - |
| **S38** | - | - | - | - | - | - |
| **S39** | - | - | - | - | - | - |
| **S40** | - | - | - | - | - | - |

**Table S1.** qPCR Ct value and OPTIMAL Tt value of the clinical samples for CMV, EBV and VZV detection. For qPCR, Samples with Ct value <40 were determined to be positive, and samples with Ct value ≥40 were determined to be negative. For OPTIMAL, Samples with Ct value <20 were determined to be positive, and samples with Ct value ≥20 were determined to be negative. “-” represents negative determination.

**Table S2.** Primers used in this study.

| Target | Name | Sequence (5'-3') |
|---|---|---|
| SARS-CoV-2 N gene | F3 | AACACAAGCTTTCGGCAG |
| | B3 | GAAATTTGGATCTTTGTCATCC |
| | FIP | TGCGGCCAATGTTTGTAATCAGCCAAG GAAATTTTGGGGAC |
| | BIP | CGCATTGGCATGGAAGTCACTTTGATG GCACCTGTGTAG |
| | LF | TTCCTTGTCTGATTAGTTC |
| | LB | ACCTTCGGGAACGTGGTT |
| EBV | F3 | TGCCCTTGCTATTCCACAAT |
| | B3 | AAGCTGCACACAGTCACC |
| | FIP | AATGGACTCCCTTAGCGGGCCCATTGA GTCGTCTCCCCT |
| | BIP | TGCTGAGGTTTTGAAGGATGCGGATAT TGCAGGTAGGAGCGG |
| | LF | GTCCAGGGGCCATTCCAA |
| | LB | TAAGGACCTTGTTATGACAAAGC |
| CMV | F3 | TGTCCGTCAAAGATGACACG |
| | B3 | GGTTATCGACATGTACCCCG |
| | FIP | TGCGCGATCTGTTCAACACCATCAACG GAATTTTAGCCAGCC |
| | BIP | ACAGGTCATCCTTGCGTTGCCGCATGG CCAAGACTAACTCG |
| | LF | ATTTTCACTACGAGGCCGGGG |
| | LB | CAGGTAAAGCTCGGCCATAGTGTT |
| VZV | F3 | TGTTGGAGAAGGGTGACCG |
| | B3 | TCGGCCCTGAACCAGTTCT |
| | FIP | TCCTACAGAGTCTCCGCAGAGCACTGG AGCCCGTTGCCT |
| | BIP | GCTTCGCCTACATGCGGAACGGACCA AAAGCTGTTGCCACC |
| | LF | TGCCAGCATGGCATACCCC |
| | LB | AGACGCTACGCTCCCCGTA |

**Table S3.** gDNAs used in this study.

| Target | Name | Sequence (5'-3') |
|---|---|---|
| SARS-CoV-2 N gene | gDNA 1 | P-TCCAGCGCTTCAGCGT-P |
| | gDNA 2 | P-TCTTCGGAATGTCGCG-P |
| CMV | gDNA 1 | P-TTCCTGCAGACAGTAA-P |
| | gDNA 2 | P-TGGCCTACCTGGGCGC-P |
| EBV | gDNA 1 | P-TTTGTCTGTTATTTCA-P |
| | gDNA 2 | P-TGGTCTTTTTACAAAC-P |
| VZV | gDNA 1 | P-TAGTGTTTTTGGGCCC-P |
| | gDNA 2 | P-TATCGACGGCTCATAG-P |

**Table S4.** Reporters used in this study.

| **Name** | **Sequence (5'-3')** |
|---|---|
| SARS-CoV-2 N gene | ROX-TCTTCAGCGTTCTTCGGAATGTCGCGCA-BHQ1 |
| CMV | HEX-CGGCGTCCTGCAGACAGTAACGGCCTACCTGG-BHQ1 |
| EBV | FAM-TGTTATTTCATGGTCTTTTTACAAACTCAT-BHQ1 |
| VZV | ROX-TTGGGCCCGGTCGTATCGACGGCTCATAGCCA-BHQ2 |

**Table S5.** Plasmids used in this study.

| Name | Description | Accession number |
|---|---|---|
| *Pf*Ago-plasmid | pET28a-*Pf*Ago | WP_011011654 |
| SARS-CoV-2-N-plasmid | pUC57-SARS-CoV-2-N gene | NC045512 |
| CMV-plasmid | pUC57-KANA-CMV UL54 | NC006273.2 |
| EBV-plasmid | pUC57-KANA-EBV EBNA | NC007605.1 |
| VZV-plasmid | pUC57-KANA-VZV ORF62 | NC001348.1 |
